# The effect of self- vs. externally generated actions on timing, duration and amplitude of BOLD response for visual feedback processing

**DOI:** 10.1101/2021.06.19.449116

**Authors:** Eleftherios Kavroulakis, Bianca M. van Kemenade, B. Ezgi Arikan, Tilo Kircher, Benjamin Straube

## Abstract

It has been widely assumed that internal forward models use efference copies to create predictions about the sensory consequences of our own actions. While these predictions had been frequently associated with reduced neural processing in sensory cortices, the timing and duration of the hemodynamic response of self-generated as opposed to externally generated movements is poorly investigated. In the present study we tested the hypothesis that predictive mechanisms for self-generated actions lead to early and shorter neural processing compared with externally generated movements. Using a first and second-order Taylor approximation in terms of the temporal (TD) and dispersion (DD) derivatives of a canonical hemodynamic response function, we investigated the timing and duration of activation for self-generated and externally generated movements using a custom-made fMRI-compatible movement device. Visual video feedback of the active and passive hand movements were presented in real time or with variable delays (0 - 417 ms). Participants had to judge, whether the feedback was delayed. We found earlier feedback processing for self-generated compared to externally generated movements in several regions including the supplementary motor area, cerebellum, subcortical structures such as the putamen and visual cortices. Shorter processing was found in areas, which show also lower blood oxygen level dependent (BOLD) amplitudes, such as the SMA, occipital and parietal cortex. Specifically, earlier activation in the putamen, of self-generated movements was associated with worse performance in detecting delays. These findings support our hypothesis, that efference copy based predictive mechanisms enable earlier processing of action feedback, as potential source for behavioral effects.

## 1. Introduction

Sensory stimuli associated with self-generated movements are perceived as less intense compared to stimuli associated with externally generated movements, and often referred as sensory suppression or attenuation (Blakemore et al., 1999). On the neural level this effect has been associated with weaker blood oxygen level dependent (BOLD) activations in somatosensory areas (Blakemore et al., 1998; Desantis and Haggard, 2016; Haggard et al., 2002). It has been suggested that this phenomenon is based on an internal forward model which uses efference copies to create predictions about the sensory consequences of our own actions (Miall and Wolpert, 1996; Sperry, 1950; von Holst and Mittelstaedt, 1950; Wolpert and Flanagan, 2001). Through the efference copy mechanism, is able to monitor if there is a match or mismatch between predicted and actual sensory feedback, in case of a match self-generated stimuli it can be predicted and is less surprising than the externally generated stimuli. On the neural level this effect has been associated with reduced neural responses related to the self-generated stimuli (Blakemore and Sirigu, 2003; Wolpert et al., 2011; Wolpert and Flanagan, 2001; Wolpert and Ghahramani, 2000). Although, sensory suppression in voluntary movements versus external generated is well established, the timing of the hemodynamic response of self-generated (active) as opposed to externally generated (passive) movements is poorly investigated. Moreover, investigating relative timing of activity between conditions and brain areas helps to understand the sequence of processing across multiple activated areas of a given contrast, to interpret amplitude differences and might help to explain behavior. Predictions need to be generated before stimulus to allow fast processing. Indeed, efference copy mechanism can explain how fast processing is possible, by taking place earlier or parallel to action execution. Thus, action related predictive mechanisms should lead to an earlier, better prepared processing of action feedback.

Over the last decades event related cortical potential studies (recorded during movement preparation) showed increased cortical activity prior to self-initiated movements and little or no early activity in movements that are externally produced (Jahanshahi et al., 1995; Papa et al., 1991) in unpredictable times (Cunnington et al., 1995). Moreover, it is believed that this movement preparatory activity initiates predominantly in supplementary motor area (SMA) in a combining positron emission tomography (PET) with recording of movement-related potentials study(Jahanshahi et al., 1995). Specifically, rostral SMA played a central role in movement preparation, while caudal SMA in motor execution (Jenkins et al., 2000). Surprisingly, very few fMRI studies have investigated the timing of hemodynamic response in self-generated as opposed to externally generated movements. One of them showed that pre-SMA was activated earlier for self-generated compared to externally triggered movement (active response to a auditory cue), reflecting involvement of this region in early processes associated with the preparation for voluntary movement (Cunnington et al., 2002). Another fMRI study examined the sequential activation of motor areas in self-initiated complex movement or in response to external auditory cues; it was shown that activation in rostral SMA occurred 0.7 s earlier than in motor cortex (MI) in the externally cued execution and 2 s earlier in the spontaneous execution movement (Weilke et al., 2001). This pointed out that timing of rostral SMA activation precedes activation in MI by different time intervals depended by the form of movement initialization. Other fMRI studies have focused in determining the temporal sequence of the hemodynamic response of different anatomical areas by calculating the latencies of different brain areas in a visuomotor experiment. They found that the timing of cortical activation begun in the visual areas followed by the SMA (preparatory phase) and finally in the motor cortex (movement initiation) (Mohamed M.A. et al., 2003). The goal of the present study is to test the hypothesis that forward model prediction lead to earlier and shorter processing of feedback from voluntary actions. In this fMRI experiment we incorporated real-time and delayed visual feedback of active and passive movements, using a custom-made fMRI-compatible movement device (Arikan et al., 2019; Van Kemenade et al., 2019) to examined i) the amplitude (HRF), timing (TD) and duration (DD) of BOLD response between self-generated and externally generated movements; ii) the latency of feedback movement onset between self-generated and externally generated movements; iii) direct associations between amplitude, timing and duration of BOLD responses with behavioural data from the delay detection task. We hypothesized that earlier processing of the upcoming visual feedback in active as opposed to passive conditions is due to predictive mechanisms which probably leads to reduce neural activity for processing of the feedback, reflected in reduced amplitude and duration of BOLD response, and it is also might correlates with reduce behavioral performance. Moreover, a possibly positive correlation between earlier processing and reduced delay detection performance for self- vs. externally generated movements, as an earlier representation of the visual motion feedback would be perceived as temporally closer to the actual movement (less delayed).

## 2. Methods

Participant’s data were taken by recent studies of our group (Arikan et al., 2019; Van Kemenade et al., 2019), which examined the naturalistic action feedback of active versus passive movement. Data acquisition and experimental design have been reported previously and are described in brief below.

### 2.1 Participants

Twenty-three right handed participants with no history of psychiatric or neurological disorders and no current use of psychoactive medications took part in the fMRI experiment. All participants had normal hearing and vision or corrected to normal vision. Right-handedness was confirmed by Edinburgh Handedness Inventory (Oldfield, 1971). The experiment was approved by the local ethics committee and performed in accordance with the Declaration of Helsinki. Three participants had to be excluded, one due to excessive head movement and two due to technical issues, so the final sample consisted of 20 participants (9 females, age = 26 ±3.24). Using a behavioral training session all participants were trained prior to scanning.

### 2.2 Equipment

A custom-made MRI-compatible passive movement device (PMD) was used for the execution of both active and passive movements (Figure 1a). The device consisted by a knob that the participant could grab and move using their wrist in active condition. The knob was moved automatically, with approximate force of 20N in the passive trials, by using compressed air (6 bar). The movement was circular from left to right and the angle between the start/ending and turning point was approximately 30^0^ (Kemenade et al., 2019; Arikan et al., 2019). There was no significant difference in movement durations between active and passive conditions. Self-generated and externally generated movements were recorded by a MRI-compatible camera (MRC High Speed, MRC Systems GmbH, Heidelberg, Germany; refresh rate: ∼4 ms). The video recordings were shown to the participants on a mirror screen. Between actual movements and camera images, variable delays (0, 83, 167, 250, 333, or 417 ms) established in previous experiments (Schmalenbach et al., 2017; Straube et al., 2017; van Kemenade et al., 2017, 2016), were inserted. Auditory stimulus (sine wave 440 Hz, 500ms duration) was presented via MR-compatible headphones (MR-Confon, Optimel, Magdeburg, Germany). Detection of movement onsets achieved by infrared LEDs, attached to the device (van Kemenade et al., 2019; Arikan et al., 2019).

**Figure 1.**
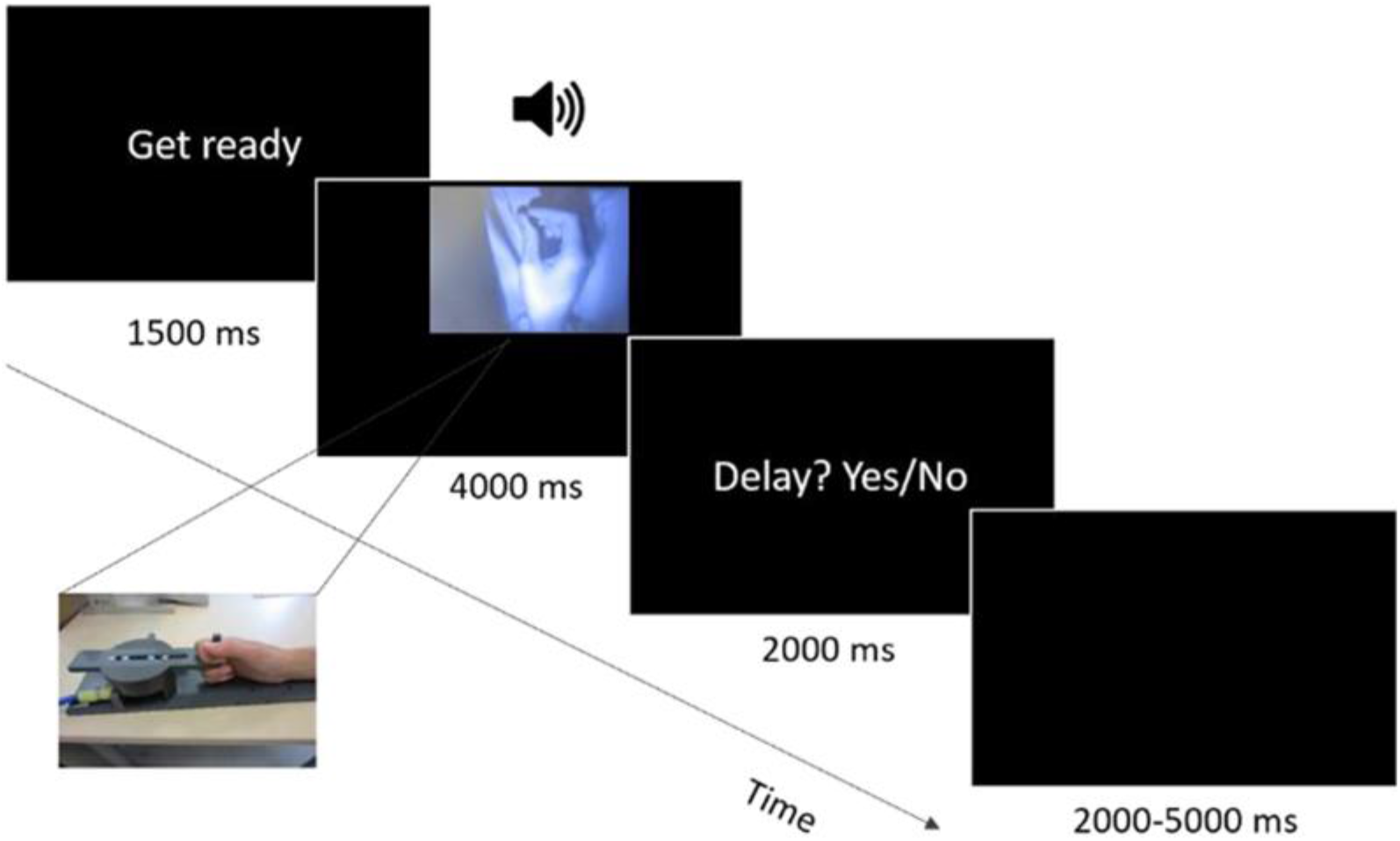
Experimental description and time frame of one trial. After the cue (“Get ready”), the camera was turned on for 4000 ms and during this time window participants had to perform a voluntary (active) or an externally generated movement (passive) based on the initial guideline. Participants observed a visual display of their movement, while their task was to judge if there was a delay or not between the actual movement and the visual feedback of the movement. During bimodal trials, an auditory stimulus was presented, coupled with the onset of the visual stimulus.

### 2.3 Task and Stimuli

The task and stimuli had been previously described (van Kemenade et al., 2019; Arikan et al., 2019) and were similar to task and/or feedback in our recent fMRI experiments (Pazen et al., 2020; Schmitter et al., 2021; Uhlmann et al., 2020). Briefly, participants performed movements with their wrist, from left to right, using the PMD during the active phase of the task. In passive condition they were instructed to hold the knob of the PMD as relaxed as possible, letting the device move their hand. Both movements were recorded by a high speed, MRI compatible camera and presented to the participants via a mirror-projection screen. Between the actual movement and the visual feedback of the movement, six delays (0, 83, 167, 250, 333, or 417 ms) were inserted and participants were asked to judge if there was a delay between them. Half of the trials consisted of a visual feedback, while the other half consisted of both visual and auditory feedback {440-Hz sine wave pure tones (500 ms)}. The paradigm consisted of five experimental runs. Each run was divided in passive and active blocks (with 24 trials each) in counterbalanced order. Each block started with written instructions, notifying participants about the movement they had to perform (active or passive). Each trial started with a cue, ‘‘Ready,’’ that lasted for 1500 ms to inform the participants that the trial was about to begin. After that, the camera was on for 4000 ms and the participants should perform the movement during that time frame in active trials, or let the PDM execute the movement for them (passive blocks). The onset of the passive trials was jittered (500 - 1500 ms). Directly afterwards, the question “Delay? Yes/No” appeared on the screen and participants had to respond, whether there was a delay or not, using their left index and middle fingers (button assignment was counterbalanced across participants) with maximum response time 2000 ms (Figure 1). In half of the trials only visual feedback (unimodal trials) was presented and in the other half both visual and auditory feedback (bimodal trials) was presented (randomized across each block). Each run had 2 trials per delay, per condition, leading to 12 unimodal and 12 bimodal trials for each block (active/passive), and thus to 48 trials per run. Every movement was recorded and monitored online to ensure agreement with instructions and for posthoc analysis of movement durations. A behavioral training session was introduced prior to scanning, in order to familiarize the participants with the task and the equipment. Specifically, participants were shown how to perform the movements using the PMD. In order to perform the movements with a constant pace, participants were trained with a metronome. The scope of that was to discard any differences between self- and externally-generated movements and any individual speed differences between participants. Afterwards, in order to familiarize themselves with the trial sequence, participants were shown one trial for each condition without delay. After that, they performed one active and one passive training run with 8 trials each (4 without delay, 4 with the maximum delay of 417ms, half of which were unimodal and half bimodal), in which they were informed about their performance (delay detection) afterwards. Finally, the participants performed 3 runs similar to the experimental run in the scanning sessions, without seeing their own hand (a curtain blocked their field of view) and were informed about their performance. Subjects with a detection rate of less than 50% at the 0 ms delay and a detection rate of at least 50% at the 417 ms delay participated to the fMRI experiment. Only one participant had to be excluded for not meeting the aforementioned criteria.

### 2.4 Image acquisition

Functional MRI data were acquired using a 3 T TIM Trio scanner (Siemens, Erlangen, Germany), using a 12-channel head-coil. A gradient echo EPI sequence was used (TR: 1650 ms, TE: 25 ms, flip angle: 70°, slice thickness: 4mm, gap: 15%, voxel size: 3 × 3 ×4.6mm). For each run, 330 volumes were obtained, each containing 34 slices covering the whole brain, acquired in descending order. Anatomical images were obtained using a T1-weighted MPRAGE sequence (TR: 1900ms, TE: 2.26ms, flip angle: 9°, slice thickness: 1mm, gap: 50%, voxel size: 1 × 1 × 1.5mm).

### 2.5 Behavioral data analysis

For each participant logistic psychometric functions were fitted to the data using Psignifit (Fründ et al. 2011). Slopes and threshold (delay at which a 50% detection rate was reached) of the psychometric functions for each condition were extracted and used for correlations with the fMRI data. For the calculation of movement durations, video recordings were used and movement onsets and offsets were extracted.

### 2.6 Preprocessing and analysis

Image processing and statistical analyses were performed in SPM12 (Statistical Parametric Mapping software, SPM: Welcome Department of Imaging Neuroscience, London, UK; available at: http://www.fil.ion.ucl.ac.uk/spm/). Realignment was applied to functional data to correct for head movement. The anatomical image of each participant was coregistered to their functional image, segmented and normalized to the standard Montreal Neurological Institute (MNI) template. The resulting parameters were then used to normalize the functional images to the MNI space (resampled to a voxel size of 2 × 2 × 2mm). Finally, the data were smoothed with an 8 ×8 × 8mm 3 full-width at half maximum kernel. A general linear model (GLM) was set up for each participant to analyze the preprocessed functional (Frinston et al.1995). The model included regressors for each condition (active unimodal, active bimodal, passive unimodal, passive bimodal) modelling the onset of the movements (active or passive) until the offset of them. Also additional regressors of the cue and the presentation of the question (until the participants answered by button press) were included. Lastly, six motion regressors of no interest were added in the final design matrix. Bimodal trials in which no tone was played or trials with no movement performed were excluded (1.4% of all trials). Hemodynamic responses were modelled by using a first and second order multivariate Taylor expansion of the canonical hemodynamic response function (HRF) (Friston et al.1998). The partial derivatives with respect to time (temporal derivative) and duration (dispersion derivative) are included in those functions. Adding this set of “basis” functions within the GLM allows estimation of the contribution of each basis function (its parameter estimate), and calculation of the mean and standard error of the best-fitting event-related response by linear combination. Specifically, inclusion of the temporal and dispersion derivative within the GLM permits differences in the latency and the duration of hemodynamic response onset to be accommodated in the model and gives the opportunity to perform statistical assumptions based on latency and duration differences between conditions (Frinston et al., 1998). Response latencies (relative to the canonical HRF) were estimated via the ratio of derivative to canonical parameter estimates (Henson et al., 2002).

On single subject level T and F-contrasts for the canonical and derivative terms were created for each regressor of the experimental conditions (active unimodal, active bimodal, passive unimodal, passive bimodal). Also an F and T contrasts were created for passive unimodal + passive bimodal > active unimodal +active bimodal. To investigate any effect of modality we contrasted bimodal against unimodal (and the opposite). We found no significant interaction of modality (bimodal vs. unimodal conditions) by movement (active vs. passive) in amplitude, timing or duration. Therefore, we used active unimodal and bimodal together as one condition (self-generated contrast), the same logic was used for passive unimodal and bimodal (external generated movements). We first examined the general effects of amplitude, TD and DD for self-generated compared to externally generated movements by using F-contrasts (for single subject analyses) and repeated-measures One-way ANOVA for second level analyses. At the single subject level, an F-contrast with three rows was specified: one with ones in the canonical HRF (amplitude) terms (one per session); ones in the temporal derivative terms (also one per session) and another with the dispersion derivative terms (again one per session).

For the second-level analyses, individual contrasts were obtained for each of the canonical and derivative terms from each subject (three contrast volumes per subject) and these volumes entered into a repeated-measures One-way ANOVA (without constant) model with one factor that has three levels (one for the canonical, second for the temporal derivative effects and third for dispersion derivative term). In this ANOVA, unequal variance across levels of factors (to account for different variances between the canonical and derivative terms) and not sphericity (to account for possible across-subject correlations between the two terms) were assumed. The following **contrasts of interest** were performed to test our hypotheses: First we tested whether active conditions were related to earlier BOLD responses by calculating a t-test active > passive regarding the effect of the temporal derivatives. Second, we tested if earlier processing is related to behavioral suppression for active vs. passive condition. Therefore, we extracted eigenvariates of the TD contrast estimates for each cluster showing the active>passive difference and performed correlation analyses 1) with the threshold differences (active-passive). Finally, we explored the effects of our movement manipulation (active/passive) on amplitude (active<passive: suppression) and duration (active<passive: suppression) of the BOLD response by comparing conditions regarding HRF and dispersion derivatives, respectively.

Family-wise error (FWE) correction at *p* < 0.05 based on Gaussian random field theory (Worsley et al., 1996; Worsley et al., 1992) was used, to correct for multiple comparisons at the whole brain level. The above approach for estimating the amplitude ignores the potential for an amplitude bias induced by a delay difference between the hemodynamic model and the data. To diminish the effect of amplitude bias due to the use of only the nonderivative portion of the model in the test for significant amplitudes, we used the approach described by Calhoun *et al*. 2004, which tested an amplitude estimate that is a function of both the nonderivative and the derivative terms of the model (Calhoun et al., 2004). In this approach, the proposed amplitude test does not suffer from delay-induced bias, comprises a more natural test for amplitude differences when using a model incorporating derivative terms, while it improves the fit of the model to the data (when compared to a model not using the derivatives term).

We combined the temporal and dispersion derivative to re-estimated the biased amplitude regressors (computation of boosted contrast maps) (Calhoun et al., 2004; Cignetti et al., 2016; Pernet, 2014) and then we used the corrected boosted contrasts at the 2^nd^ level (as described above). Specifically, we estimate the time-to-peak of the BOLD response voxel-wise, then a mask of the voxels whose responses peaking was within the specified temporal range (full range of the basis set) was created, then the parameter estimates were replaced with their boosted equivalents for voxels within the mask and finally, the contrast of interest were re-estimated. The code (spmup_hrf_boost.m) is available at the GitHub archive: https://github.com/CPernet/spmup/blob/master/spmup_hrf_boost.m). Finally, we calculated the latency differences between passive and active conditions within several brain regions, via estimated the ratio of temporal derivative to amplitude parameter estimates and transformed this ratio for each voxel with a sigmoid logistic function (Henson et al., 2002).

## 3. Results

### 3.1. Behavioral analysis

Behavioral results had been already reported (van Kemenade et al., 2019), and are here especially important for the correlation with neural data. In brief, slopes and thresholds (delay at which a 50% detection rate was reached) of the psychometric functions for each condition were used from the behavioral data. Repeated-measures ANOVAs were applied on the slopes and the thresholds with the factors Modality (unimodal/bimodal) and Action (active/passive). Significant main effect of Action (F(1,19) = 6.779, P = 0.017, η^2^ p = 0.263), with lower threshold and thus better performance in passive trials was observed. No significant main effect of Modality (F(1,19) = 1.091, P = 0.309, η^2^ p = 0.054) or interaction with Action (F(1,19) = 1.345, P =0.261, η^2^ p = 0.066). Also, no significant main effects using the slopes were found, neither effect of Action (F(1,19) = 1.156, P = 0.296, η^2^ p = 0.057 or of Modality (F(1,19) = 1.971, P = 0.176, η^2^ p = 0.094) or interaction ((F(1,19) =1.283, P = 0.271, η^2^ p = 0.063). Also, there was no significant differences (P = 0.234, d = 0.340) in duration between condition, so any possible differences between them cannot be attributed in that (van Kemenade et al., 2019).

### 3.2 Timing in externally generated versus self-generated movements

By contrasting externally with self-generated movements, we revealed an active > passive difference for the effect of the TD (table 1), indicating earlier hemodynamic responses in active versus passive conditions, in bilateral middle occipital gyrus (MOG) and superior occipital gyrus (SOG), in subcortical regions including bilateral putamen, thalamus and caudate nucleus. Also, effects indicating earlier BOLD responses for active conditions were found in bilateral supplementary motor area (SMA), superior frontal gyrus (SFG), insula, and in superior parietal lobule (SPL) and inferior parietal lobule (IPL). Moreover, the contrast revealed effects in parietal, temporal, pre-motor and cerebellar areas (cerebellum crus I, II and lobule VI) and specifically were activated earlier in active compared to passive condition (Figure 2). Anatomical locations and detailed description of those clusters are shown in Table 1.

**Table 1.** Anatomical locations of earlier activations (TD) for self-generated (active) versus externally-generated (passive) movements. L: Left; R: right; MOG: Middle Occipital Gyrus; SOG: Superior Occipital Gyrus; SFG: Superior Frontal Gyrus; MFG: Middle Frontal Gyrus; DLPFC: Dorsolateral Prefrontal Cortex; MTG: Middle Temporal Gyrus; ACC: Anterior Cingulate Cortex; IPL: Inferior Parietal Lobule; SPL: Superior Parietal Lobule; SMA: Supplementary Motor Area (pFWE < 0.05)

|  |  | Active-Passive (TD) |  |  |  | Cluster size |
| --- | --- | --- | --- | --- | --- | --- |
| Brain area | Hemisphere | x | y | z | T |  |
| Precuneus | L | -12 | -68 | 58 | 5.27 | 14 |
| Rolandic_Operculum | L | -50 | 2 | 14 | 5.24 | 106 |
| Thalamus | L | -10 | -14 | 10 | 7.29 | 261 |
| Lingual | L | -2 | -70 | 6 | 5.92 | 247 |
| Calcarine | L | -4 | -92 | 6 | 8.09 | 657 |
| MOG | L | -32 | -92 | 6 | 8.49 | 586 |
| MOG | R | 42 | -84 | 4 | 9.48 | 585 |
| SOG | L | -12 | -98 | 4 | 6.69 | 589 |
| SOG | R | 18 | -98 | 6 | 7.38 | 490 |
| Thalamus | R | 20 | -20 | 2 | 7.13 | 176 |
| SFG | L | -22 | 50 | 8 | 6.46 | 108 |
| SFG | R | 26 | 56 | 8 | 5.54 | 155 |
| MFG | L | -40 | 42 | 24 | 7.19 | 758 |
| MFG | R | 36 | 58 | 2 | 6.55 | 1673 |
| DLPFC | R | 52 | 12 | 36 | 6.54 | 1673 |
| Insula | L | -48 | 18 | -10 | 5.15 | 16 |
| Insula | R | 50 | 16 | 2 | 6.81 | 91 |
| Putamen | L | -24 | 4 | -8 | 8.73 | 119 |
| Putamen | R | 30 | 2 | 0 | 7.83 | 118 |
| Caudate | L | -14 | 10 | 16 | 5.4 | 14 |
| Caudate | R | 16 | 12 | 8 | 5.33 | 24 |
| ACC | L | -4 | 42 | 16 | 5.26 | 2 |
| IPL | L | -28 | -50 | 38 | 5.28 | 4 |
| IPL | R | 32 | -48 | 48 | 5.47 | 11 |
| Postcentr | L | -52 | -8 | 50 | 7.27 | 74 |
| Precentral | L | -50 | -6 | 50 | 7.04 | 239 |
| Precentral | R | 40 | -16 | 56 | 7.14 | 187 |
| Cerebel VI | R | 40 | -60 | -26 | 7.75 | 596 |
| Cerebel VI | L | -36 | -64 | -24 | 6.96 | 384 |
| Cereb_crus_I | L | -38 | -64 | -26 | 6.05 | 7670 |
| Cereb_crus_I | R | 40 | -60 | -26 | 7.75 | 596 |
| Cereb_crus_II | L | 0 | -82 | 28 | 5.23 | 5 |
| SPL | L | -20 | -56 | 48 | 5.23 | 43 |
| SPL | R | 26 | -66 | 32 | 5.13 | 41 |
| SMA | L | -4 | 12 | 46 | 7.95 | 232 |
| SMA | R | 12 | 14 | 44 | 6.98 | 172 |
| Cerebel_Crus_II | R | 2 | -82 | -28 | 5.23 | 5 |

**Fig 2.**
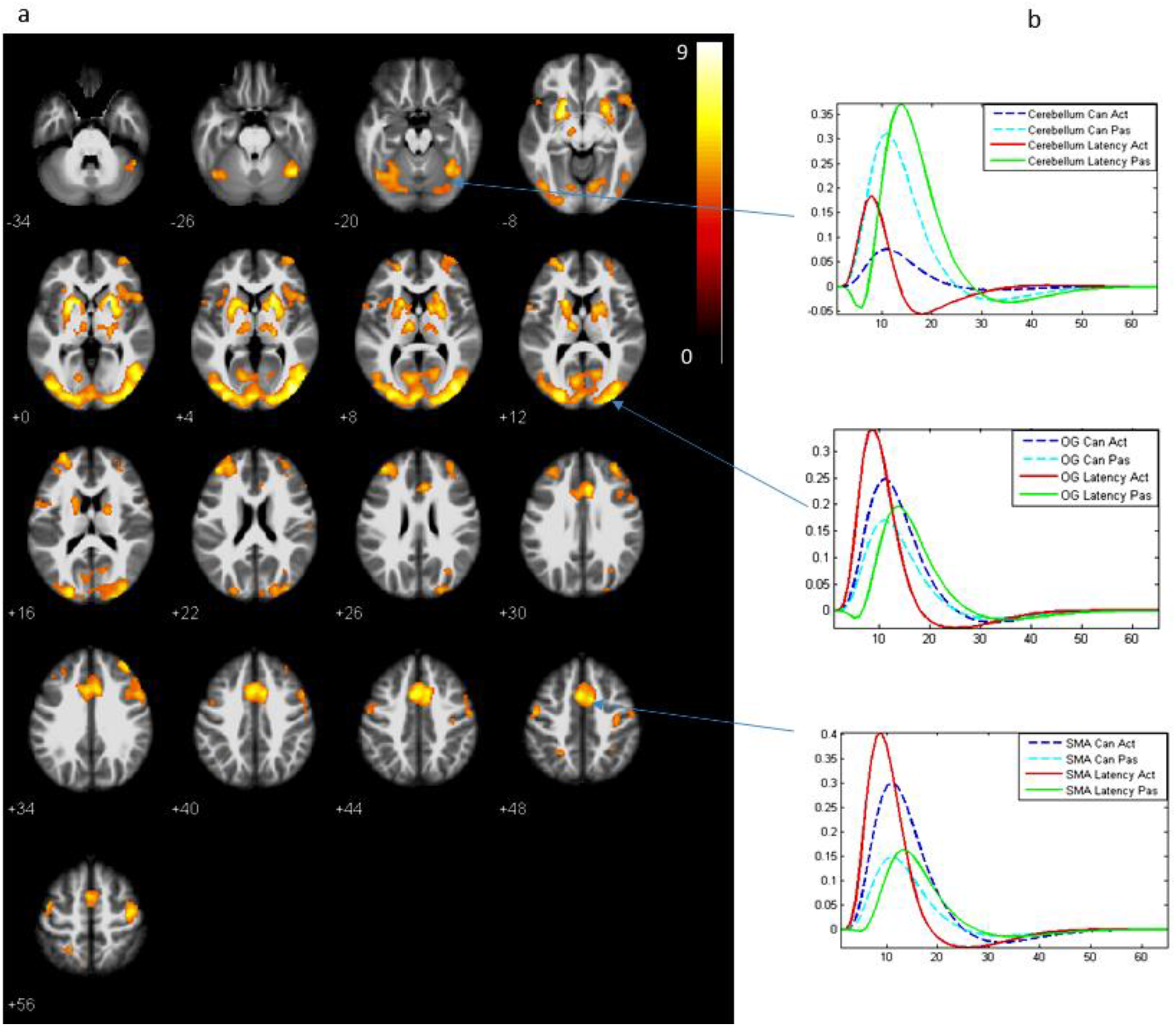
a. Earlier hemodynamic response in active versus passive conditions in cortical and sub-cortical areas. b. Latencies for active (red curve) and passive (green curve) conditions, indicated that Cerebellum, OG (Occipital gyrus) and supplementary motor area (SMA) activate earlier in active condition that in passive (the same trend also found in the other brain areas). Blue curve corresponds to the amplitude of the canonical HRF for activate and light blue for passive.

#### 3.2.1 Timing and delay detection performance

Positive correlations were found between the timing of the hemodynamic response and behavioral threshold values in the left putamen (*r* = 0.4782*) in passive minus active conditions (Figure 3a). This indicated that sensory suppression (negative values for passive-active) was correlated with earlier activation (negative values for pas-act) in active vs passive conditions.

**Figure 3.**
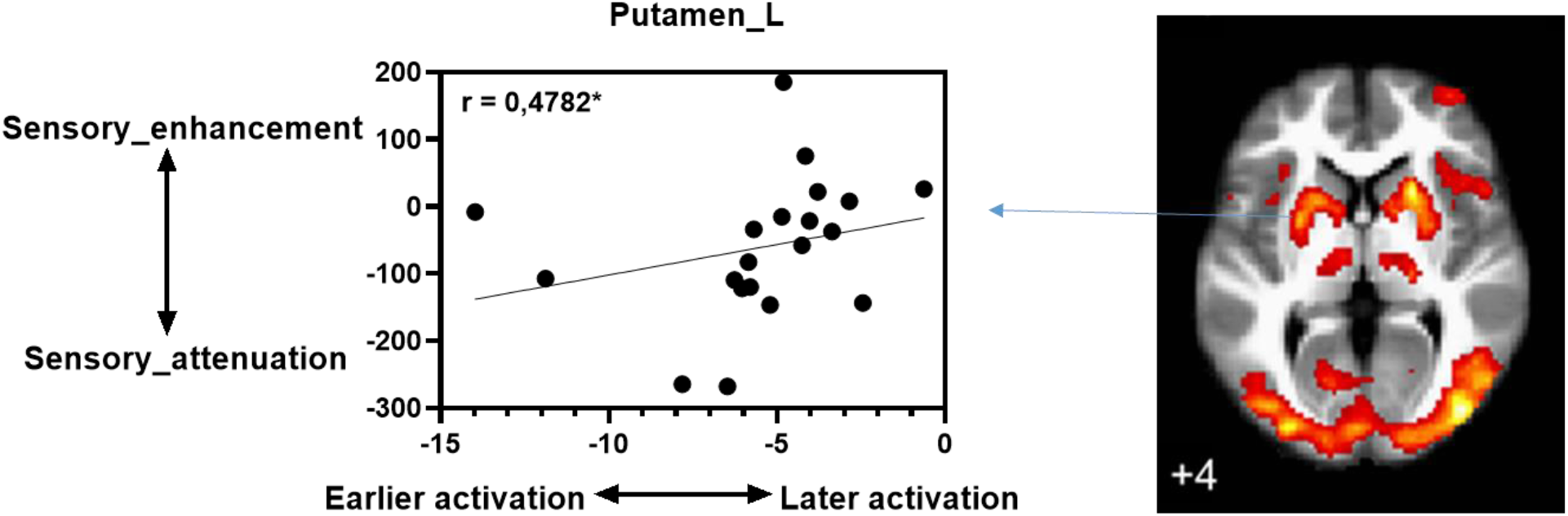
Positive correlation between the timing of the hemodynamic response and behavioral threshold values in the left putamen (r = 0.4782*), indicates that sensory suppression was correlated with earlier activation in active vs passive conditions.

**Figure 4.**
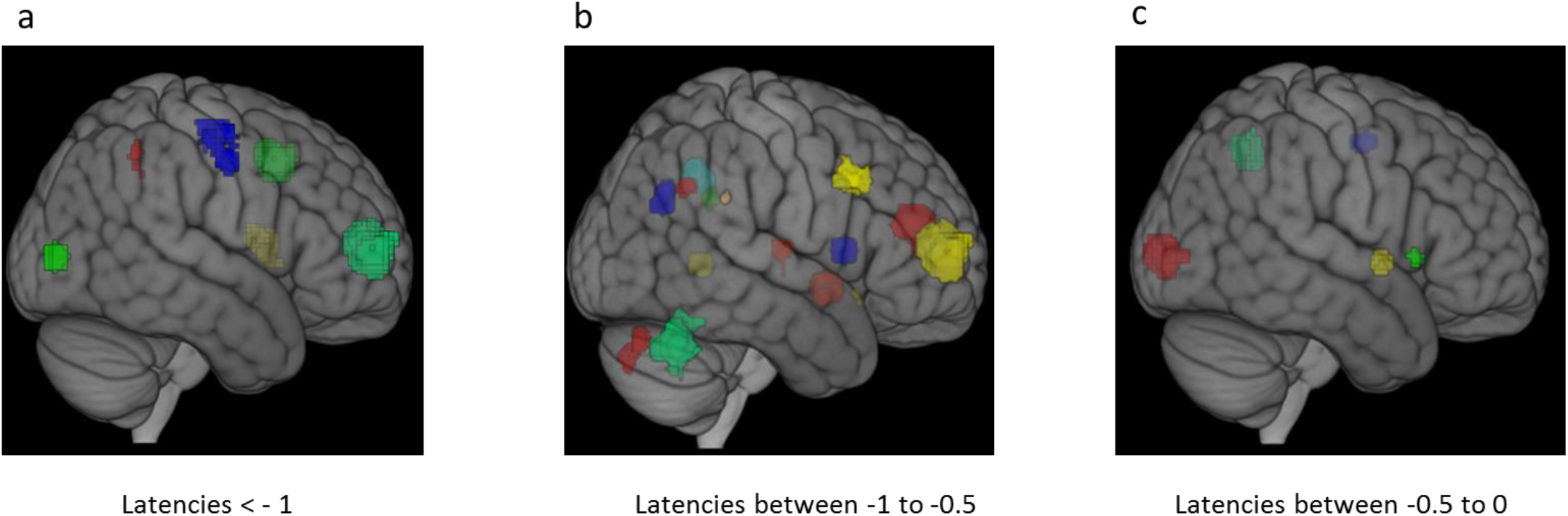
3D rendering of latency clusters during self-generated (active) as opposed to externally (passive) generated movements within several brain regions in three different time periods (in seconds (s)). a. < -1 s. b. -1 s to -0.5 s and c. -0.5 s to 0. Negative values indicate earlier activation for active versus passive conditions

#### 3.2.2 Calculation of latency onsets in self-generated as opposed to externally generated movements

Response latencies (relative to the canonical HRF) were estimated via the ratio of temporal derivative to canonical parameter estimates (Henson et al., 2002). Specifically, we calculated the feedback latency onset differences between self-generated and externally generated movements within several brain regions (Table 2). Activity raised earlier in SMA, occipital cortex, inferior and superior parietal lobule and superior frontal cortex, followed by subcortical areas, middle temporal gyrus, anterior cingulate cortex and cerebellum in active as opposed to passive condition.

**Table 2.** Latency onsets of peak activation clusters during self-generated (active) as opposed to externally (passive) generated movements within several brain regions in three different time periods from -1 to -0.5, -0.5 to 0 and <-1. Negative values indicate earlier activation for active versus passive conditions. L: Left; R: right; MOG: Middle Occipital Gyrus; SOG: Superior Occipital Gyrus; SFG: Superior Frontal Gyrus; MTG: Middle Temporal Gyrus; ACC: Anterior Cingulate Cortex; IPL: Inferior Parietal Lobule; SPL: Superior Parietal Lobule; SMA: Supplementary Motor Area; DLPFC: Dorsolateral Prefrontal Cortex (pFWE < 0.05).

| Latency onset < -1 | Latency onset between<br>-1 to -0.5 | Latency onset<br>between -0.5 to<br>0 |
| --- | --- | --- |
| SMA<br>Rolandic_Operc_L<br>MOG_R<br>SFG_R<br>IPL_R<br>Precentral_R | Ang_L<br>Precuneus_R<br>Precuneus_L<br>Supramarginal_L<br>Lingual_L<br>Thalamus_L<br>SFG_L<br>Insula_L<br>Putamen_L<br>MTG_L<br>Caudate_R<br>Caudate_L<br>ACC<br>IPL_L<br>Supramarginal_R<br>Cerebellum_R<br>Cerebellum_Crus_II<br>DLPFC<br>SPL_R | Calcarine_L<br>Insula_R<br>Putamen_R<br>Postcentral_L<br>SPL_L |

### 3.3 Amplitude and duration of BOLD responses in externally generated versus self-generated movements

Amplitude and duration of BOLD response, reflected in the effects of the HRF and dispersion derivative, respectively, were also examined in externally versus self-generated movements. Suppression (negative values for passive>active) was observed in bilateral SMA, angular, lingual gyrus, middle temporal gyrus (MTG), precuneus, insula, inferior parietal lobule (IPL) and superior frontal gyrus (SFG). Also in the left caudate, in supramarginal and in the right cerebellum and precentral gyrus (Table 3). Significant shorter processing time (negative values for passive>active) in active versus passive conditions was observed in MTG, caudate bilaterally and in right SFG, inferior frontal gyrus (IFG), inferior temporal gyrus (ITG), cerebellum crus II and in calcarine cortex (Table 4).

**Table 3.** Anatomical locations of peak activations in externally (passive) versus self-generated (active) movements. L: Left; R: right; SFG: Superior Frontal Gyrus; MFG: Middle Frontal Gyrus; Supramarg: Supramarginal; MTG: Middle Temporal Gyrus; ITG: Inferior Temporal Gyrus ACC: Anterior Cingulate Cortex; IPL: Inferior Parietal Lobule; SPL: Superior Parietal Lobule; SMA: Supplementary Motor Area; Cerebel: Cerebellum; (pFWE < 0.05)

|  |  | Passive-Active (Amplitude) |  |  |  |  |
| --- | --- | --- | --- | --- | --- | --- |
| Brain area | Hemisphere | x | y | z | T | Cluster size |
| Supramarg | L | -46 | -44 | 30 | 5.92 | 40 |
| Angular | L | -36 | -56 | 34 | 5.65 | 90 |
| Angular | R | 58 | -58 | 30 | 5.16 | 21 |
| Precuneus | R | 4 | -56 | 34 | 5.38 | 223 |
| Precuneus | L | -4 | -56 | 34 | 5.95 | 223 |
| SFG | L | -22 | 50 | 8 | 5.92 | 80 |
| SFG | R | 26 | 56 | 8 | 6.88 | 441 |
| Insula | L | -30 | 20 | -6 | 7.16 | 166 |
| Insula | R | 30 | 28 | -6 | 5.39 | 36 |
| MTG | L | -48 | -50 | 2 | 5.78 | 343 |
| MTG | R | 44 | -60 | 2 | 5.90 | 1123 |
| Caudate | L | -16 | 8 | 12 | 5.13 | 4 |
| ACC | L | -2 | 16 | 22 | 5.2 | 1428 |
| ACC | R | 4 | 40 | 24 | 5.67 | 1428 |
| IPL | L | -46 | -48 | 36 | 5.33 | 40 |
| IPL | R | 42 | -54 | 36 | 5.27 | 21 |
| Cerebel VI | R | 38 | -66 | -30 | 5.68 | 596 |
| Cerebel_Crus_II | R | 6 | -80 | -34 | 5.19 | 7 |
| SPL | L | -20 | -56 | 48 | 5.31 | 43 |
| SMA | L | -4 | 12 | 46 | 5.65 | 232 |
| SMA | R | 12 | 14 | 44 | 5.20 | 172 |
| ITG | R | 46 | -54 | -12 | 5.3 | 1123 |
| Lingual | L | -10 | -72 | 0 | 5.26 | 4 |
| Lingual | R | 16 | -74 | 0 | 5.73 | 1123 |
| MFG | L | -34 | 50 | 0 | 5.2 | 228 |
| MFG | R | 34 | 56 | 0 | 6.26 | 441 |

**Table 4.** Anatomical locations of shorter BOLD duration (DD) during self-generated (active) versus externally (passive) generated movements. L: Left; R: right; SFG: Superior Frontal Gyrus; IFG: Inferior Frontal Gyrus; ITG: Inferior Temporal Gyrus; MTG: Middle Temporal Gyrus; (pFWE < 0.05)

|  |  | Active-Passive (DD) |  |  |  |  |
| --- | --- | --- | --- | --- | --- | --- |
| Brain area | Hemisphere | x | y | z | T | Cluster size |
| SFG | R | 24 | 52 | 10 | 5.39 | 62 |
| Calcarine | R | 30 | -68 | 10 | 5.8 | 38 |
| IFG | R | -34 | 36 | -2 | 5.23 | 20 |
| MTG | L | -46 | -14 | -18 | 5.28 | 10 |
| MTG | R | 46 | -48 | -4 | 5.66 | 9 |
| Caudate | L | -18 | 12 | 14 | 5.24 | 2 |
| Caudate | R | 18 | 20 | 10 | 5.75 | 22 |
| Cerebellum_Crus_II | R | 4 | -84 | -38 | 5.17 | 3 |
| ITG | R | 48 | -46 | -6 | 5.39 | 9 |

#### 3.3.1 Amplitude, BOLD duration and delay detection performance

Exploration analysis (F contrast), revealed negative correlations between amplitude and behavioral threshold values in the left middle occipital (*r =* -0.4735*), lingual gyrus (*r =* - 0.5316*), SFG (*r =* -0.4674*), putamen (*r =* -0.4743*), rolandic operculum (*r =* - 0.4571*), precentral gyrus (*r =* -0.6602**) and in the right precentral (*r =* -0.5080*), and SOG (*r =* 0.4909*) in passive minus active conditions (Supplementary Figure 6). This indicated that sensory suppression (negative values for passive-active) was correlated with BOLD suppression (positive values for passive-active) in active versus passive conditions. Contradictory, positive correlations were observed between dispersion derivative differences (reflecting BOLD duration or processing time differences) and delay detection performance. Particularly, in left SFG (*r =* 0.4676*), putamen (*r =* 0.4416*), caudate nucleus (0.4978*), MTG (*r =* 0.4508*), cerebellum crus II (0.5634**), rolandic operculum (*r =* 0.5128) and bilateral SOG (*r =* 0.4541-Left, *r =* 0.5008) (Supplementary Figure 7). Shorter processing time (negative values in passive-active) was correlated with sensory suppression (negative values for passive-active) in self-generated movements in comparison to with the externally generated ones.

## 4. Discussion

In the present study we used a delayed visual feedback of self-generated and externally generated movements, using a custom-made fMRI-compatible device (1) to examine whether forward model predictions lead to earlier processing of feedback from self-generated actions, (2) to identify the sequencing of the neural activity of different brain areas during voluntary actions and (3) to estimate hemodynamic amplitude and width differences between self-generated as opposed to externally generated movements. We found, earlier hemodynamic response in active as opposed to passive conditions in cortical, cerebellar and subcortical brain regions. Also, earlier activity in left putamen was correlated with sensory suppression in self-generated versus externally generated movements. Moreover, that earlier BOLD response started in SMA, in middle occipital gyrus, in parietal and frontal cortex followed by cerebellum and subcortical areas like caudate, putamen and thalamus. Finally, we found suppression and shorter BOLD duration in visual, sensory, premotor and subcortical areas in active versus passive conditions. Our findings indicate, that predictive mechanisms enable the brain to start earlier with the processing of upcoming visual movement feedback in self-generated as opposed to externally generated movements. This predictive processing seems to reduce the neural resources spent for feedback processing reflected in reduced amplitude and duration of BOLD response. However, in line with the cancelation account, the predictive processing is also correlated with reduced behavioral performance.

### 4.1. Earlier processing of self-generated movement feedback

By contrasting self-generated versus externally generated movements, elicited earlier cortical activations in somatosensory, in visual, in frontal areas and also in subcortical areas and cerebellum. Specifically, we found that SMA is earlier activated for feedback processing of self-generated movements versus feedback of externally generated movements. Previous, studies reporting that pre-SMA activation occurred significantly earlier for the self-initiated movements, compared with externally generated ones (Cunnington et al., 2002). The aforementioned study concluded that preparatory activity in pre-SMA in readiness for action reflects involvement in early processes related with the preparation for voluntary movements (Cunnington et al., 2002). Other fMRI studies have shown increased activation of the SMA during early stages of movement preparation prior to movement onset (Lee et al., 1999; Richter et al., 1997; Wildgruber et al., 1997). Furthermore, in another study SMA began to be activated several seconds before self-generated movements (Sakata et al., 2017). EEG studies have revealed a negative potential signal, referred as readiness potential (RP), which starts up to one to two seconds before self-initiated movements (Kornhuber and Deecke, 1965) and its origin is the SMA (Shibasaki and Hallett, 2006). Studies using high-density EEG recording methods found an early preparatory activity prior to self-initiated voluntary movement onset in the SMA (Ball et al., 1999; Cui and Deecke, 1999). Similar reports have suggested that the SMA is a key structure for controlling self-initiated movements (Jäncke et al., 2000). Moreover, it was suggested that SMA comprises one component of a system crucial for the timing of internally triggered movements (Rao, 1997). Thus, in our study a better preparation for feedback processing, due to forward model predictions, in active compared to passive conditions can explain earlier BOLD responses in the SMA.

However, it is unlikely that this early activity in SMA is the main and potentially the only origin of self-initiated movements (Eccles, 1982) and movement feedback from self-generated movements, which in turn might be also one of the source of efference copy based predictions. In our study we reported earlier activation in additional cortical areas other than the SMA for feedback processing of self-generated movements as opposed to externally generated movements. Specifically, earlier activation in IPL, in SPL and in SFG for self-generated movements agrees with the fact of early activation in frontal and parietal cortices, together with the SMA, predicting the outcome of free decision before the participant registers awareness of those decisions (Soon et al., 2008). Another study demonstrated that significant early activation was observed in the right frontoparietal (IFG and IPL) areas during movement performed when the participant felt the urge to do it compared to movements performed in response to a visual stimulus (Sakata et al., 2017).

We also observed earlier activation in visual and auditory cortices in precuneus and in insula in visual feedback of self- as opposed to externally generated movements. A recent study reported that these areas were significant earlier activated in self-initiated movements compared to externally triggered movements (Sakata et al., 2017). Our data suggest that this preparatory activity probably prepares the brain not only for execution of the movement, but also for monitoring and processing of the expected visual consequences.

Activation in the precuneus several seconds before movement onset led to predicted decisions during a free-decision task (Soon et al., 2013, 2008). Insula has been suggested to evaluate the outcomes of intentional action decisions (Brass and Haggard, 2010). These cortical areas probably contribute to the generation of self-initiated movements concerning advance stages of neural processing that connect neural activities in sensory cortices and SMA (Sakata et al., 2017) as potential basis for either optimal feedback processing or cancelation of self-generated inputs to free resources for processing of unexpected events.

Moreover, earlier activation was observed in, middle temporal gyrus (MTG) and left rolandic operculum. Especially, the MTG is relevant for processing and awareness of temporal discrepancies in action feedback monitoring and it probably has a role with cerebellum in the transmission of information about sensory mismatches in self-generated actions (Van Kemenade et al., 2019). However, for the aforementioned areas, no previous studies have demonstrated so far early activation during feedback processing. This might be explained by the fact that our study design is novel compared to the above mentioned studies, because we compared the feedback of pure passive movements, using the PMD, with active movements (movements performed from the participant in response to a visual or auditory stimulus) and not actions which both self-initiated and externally generated performed by the subjects with the only difference is just the cuing.

Also, we found earlier activation in cerebellum for feedback processing of self- as opposed to externally generated movements. It has been proposed that cerebellum is a comparator area (Blakemore et al., 2001) specific to self-generated actions (Van Kemenade et al., 2019) which processes sensory mismatches based on efference copy mechanisms. Earlier activity in cerebellum in voluntary actions, can be explained by the fact that a comparator area it need to build up the predicted representation by comparing the incoming sensory information with the predicted ones. An animal study has reported that cerebellum has a central role during movement preparation (Sasaki et al., 1979). Deep brain stimulation studies in patients with cerebellar stroke (Gerloff et al., 1996), with progressive myoclonic ataxia (Shibasaki et al., 1986) and with infarct of the mesial tegmentum of the midbrain (Ikeda et al., 1994) showed that cerebellar circuit enrolls in movement preparation, making it a suitable region for predictive perception of movement feedback. Moreover, an fMRI study reported that bilateral cerebellar activation precedes finger movements (Cui et al., 2000). Finally, cerebellum has a supplementary role in planning related to detection and adaptation of visuomotor errors (Elsinger et al., 2006), processes necessary to solve our applied action-feedback delay detection task

Finally, we found, sub-cortical early preparatory activity before movement feedback onset of self-generated movements as opposed to externally generated movements. Specifically, earlier activation was observed in caudate nucleus, putamen and thalamus. Previous fMRI studies, have mainly focused before movement initiation period and reported that basal ganglia prepare actions before movement onset, and suggested that striatum is involved in advance planning of internal generated actions (Elsinger et al., 2006). Positron emission tomography (PET) studies has shown, increased cerebral blood flow (CBF) in thalamus, basal ganglia and cerebellum before voluntary movements (Deiber et al., 1996; Jahanshahi et al., 1995; Wessel et al., 1997). A deep brain stimulation study showed that the cortico-basal ganglia-thalamocortical network contributes in preparation of self-generated wrist extension compared to passive (manually extended by a third person) wrist extension (Paradiso et al., 2004). Moreover, studies using single cell recordings have found activation in putamen and caudate nucleus prior to execution of self-initiated movements (Romo and Schultz, 1992; Schultz and Romo, 1992). Putamen has a significant role in the self-initiated non-routine actions (Monchi et al., 2006). Higher activation was observed in putamen in self-generated compared to externally triggered actions (Cunnington et al., 2002; Jahanshahi et al., 1995; Jenkins et al., 2000; Wiese et al., 2004), as those require more planning (François-Brosseau et al., 2009). Thus, subcortical brain regions are related to movement planning and may therefore also be related to preparation for the processing of visual movement feedback from self-generated-movements, as indicated by our results. In line with this assumption, increased activation in putamen for positive feedback has been linked with a range of different tasks including motion prediction (Ullsperger, 2006) and timing estimation (Nieuwenhuis et al., 2005). Activation in left putamen, comparing target appearance to locations predicted with 50% probability versus locations predicted with 100% probability, reflects an internally generated prediction error signal when prediction has some degree of uncertainty (Sommer and Pollmann, 2016). In our model less predictable visual feedback in passive as opposed to active movements, reflects higher degree of uncertainty and prediction error signals. Caudate nucleus is activated when planning novel responses or self-initiated non-routine movements (Monchi et al., 2006). This activation often occurs in synchrony with dorsolateral prefrontal cortex (François-Brosseau et al., 2009) consisting a ‘cognitive’ corticostriatal loop (Alexander et al., 1986), which is linked with the execution of complex planning tasks (Owen, 1997). Overall, basal ganglia have a central role in regulating parts of movement initiation such as timing and selection of future movements (Dudman and Krakauer, 2016; Redgrave et al., 1999; Thura and Cisek, 2017), and are also involved in predicting action feedback or outcome (Doya, 1999; Fermin et al., 2016). In line with this assumption, we found timing, duration and amplitude of the putamen to be correlated to action-feedback delay detection performance (see Figure. 3 and Supplementary Figures 6, 7). Thus, for the first time we were able to connect this subcortical activation with action outcome monitoring, suggesting a potential role in generating predictions and preparing the brain for the likely upcoming visual information. The effort for this process seem to be low, as indicated by reduced BOLD response amplitude and duration, but occurs earlier, as indicated by the negative difference in the effect of the TD between passive and active conditions. By contrast for passive conditions less predictable visual feedback might be responsible for increased BOLD response amplitude and duration as well as later processing which probably reflects prediction error signals. Our findings suggest that also other regions such as the thalamus might contribute to predictive processing. We found that the thalamus is earlier activated in active versus passive movements. Increased thalamic neuronal firing rate was also observed before voluntary movements (MacMillan et al., 2004). Additionally, decreased neuronal firing rate of thalamus VL neurons was found before and during voluntary movements in patients with Parkinson’s disease and writer’s cramp syndrome (Crowell et al., 1968). Moreover, it has been shown that inhibition of thalamostriatal projections disturbs action initiation (Díaz-Hernández et al., 2018). All the aforementioned studies considered timepoints before movement initiation, while our study was focused in the perception of visual feedback of self-generated movements and showed for the first time earlier activation in thalamus related with movement feedback of active versus passive movements.

Taking together, our results concerning earlier activation in frontal areas, cerebellum and striatum for feedback processing of self-generated movements versus externally generated ones are in agreement with the fact that caudate possibly evaluates the quality of predictions occuring in the DLPFC-cerebellum circuit, while anterior putamen and ventral striatum may evaluate the goodness of the executed actions based on sensory feedback (Fermin et al., 2016). Activation in dorsolateral prefrontal cortex (DLPFC) has been linked with sequential action planning (Doya, 1999), forward planning and prediction of an action’s future outcome (Daw et al., 2005; Dayan, 2009; Gläscher and O’Doherty, 2010; Warner et al., 2013; Wunderlich et al., 2012). Specifically, an internal model starting from the cerebellum predicts the outcome from a hypothetical action. The prefrontal cortex keeps that in working memory and the striatum approves or rejects the action, based on the estimation of the predicted outcome (Doya, 1999; Fermin et al., 2016). The neural circuit consisting of cerebellum, prefrontal cortex and striatum is likely involved in action planning or mental simulation (Fermin et al., 2016) and predictive movement feedback processing.

In summary, feedback of self-generated movements is processed earlier than feedback of externally generated movements in a distributed cortical, cerebellar and sub-cortical network, consisting of motor, sensory, cerebellum and basal ganglia cortices probably due to efference copy based forward model predictions

### 4.2 Timing and sequencing of neural activity in self-generated movements compared to externally generated movements

Timing and sequencing of neural activity are vital both for planning and execution of movements; however the neural mechanisms responsible for the timekeeping processes are not well investigated. If we assume that different brain regions contribute different functions or resources to a specific task, it is likely that distinct brain regions are not activated completely simultaneously, even when identified with the same contrast. Latency onsets were calculated for different brain areas for feedback processing of self-generated movements versus externally generated movements (Table 2). For feedback processing of self-generated movements activity started in SMA 1 second earlier, as opposed to externally generated movements. Thus, processing within the SMA might even start before movement was executed and up to 1.43 seconds before movement of the visual feedback (considering the longest delay of 434ms). This is compatible with an fMRI study reported that peak latency of the hemodynamic response within the SMA was 1.48 s earlier for self-generated than for externally cued actions (Cunnington et al., 2002). Different electrophysiological studies reported that increased motor preparatory activity started 1 to 2 seconds prior to self-generated voluntary movements (Deecke et al., 1969; Deecke and Kornhuber, 1978). This preparatory activity starting before movement and feedback processing might represent a Bereitschaftpotential which is an early component with a slowly increasing negative potential starting 1 to 2 seconds before movement and its origin is the SMA (Deecke and Kornhuber, 1978). Earlier activity in ACC started around 1 second before movement initiation in self-generated movements as opposed to externally generated movements, which is close in agreement with a high density EEG recording study reporting that activity started in the anterior cingulate cortex 2.5 seconds, followed by pre-SMA 2.2 seconds prior self-generated movement onset (Ball et al., 1999). Additionally, fMRI studies have been used to examine latency differences between self-generated and externally trigger (cued) movements. Significant timing of activation difference in peak latency of the fitted hemodynamic response was found between SMA and sensorimotor cortex in self-generated movements (1.20 s). However, in externally generated movements (0.25 s) either a trend or no difference was observed (Cunnington et al., 2002). Another study showed that pre-SMA activity preceded primary motor cortex activity by 2 seconds for self-generated movements and by 0.7 seconds for externally-cued movements (Weilke et al., 2001). We also, found that activation in MOG started 1.25 seconds earlier for self-generated compared to externally generated movements, indicating early preparation for the processing of the upcoming visual movement feedback in active conditions. This is in agreement with a visuomotor study, in which the sequencing of cortical activation started in occipital cortices, followed by the SMA and lastly by the motor cortex (Mohamed M.A. et al., 2003). Moreover, activation in cerebellum, in left putamen and in caudate started 900 ms, 1 sec and 700 ms respectively, prior to feedback of self-generated movements. This is close in agreement with studies reporting that pre-movement activity during self-generated actions reverberates through cortex-basal-ganglia-thalamocortical loops with latencies of 200-250 ms (Klaus and Plenz, 2016; Oldenburg and Sabatini, 2015). Furthermore, similar results were found in other studies reporting increased neuronal activity in cerebellum and basal ganglia 300 ms prior to movement onset for self-generated limb movements (Van Donkelaar et al., 1999).

In summary, by calculating latency-differences between active and passive conditions we were able to explore the potential processing hierarchy for voluntary actions, starting with activity in the occipital structures, via SMA, Cerebellum and followed by an involvement of more distributed cortex-basal-ganglia-thalamocortical loops. Even though the latencies are just estimates and depend on many factors such as model fit, cluster size or size of a given brain region, this pattern is in line with the literature and suggests that predictive processes related to voluntary (compare to passive) movements affect the timing of the complete assemble of cerebella-basal-ganglia-thalamocortical regions and might even start with a visual representation in the occipital lobe.

### 4.3 Suppression and shorter processing of self-generated movement feedback

We found significantly BOLD amplitude suppression in self-generated versus externally generated movements in different brain regions (passive>active). Namely, in bilateral SMA, anterior cingulate cortex (ACC), precuneus, angular gyrus, insula, thalamus, middle temporal gyrus (MTG), lingual gyrus, middle frontal gyrus (MFG), inferior parietal lobule (IPL) and superior frontal gyrus (SFG). Also, in left caudate, supramarginal, SPL, calcarine and in right cerebellum VI, inferior temporal gyrus (ITG) and cerebellum crus II. These findings are consistent with our previous study, where only canonical HRF and no basis functions had been used for analyses (Arikan et al., 2019). Also, we found significantly shorter processing time (DD) in active as opposed to passive condition in bilateral caudate, MTG and in right SFG, inferior frontal gyrus (IFG), in calcarine gyrus, in ITG and in the right cerebellum crus II.

To our knowledge, our study is the first to explore BOLD processing time differences in self-generated versus externally generated movements. Estimation of hemodynamic delay and width, in addition to amplitude, provides a more detailed and quantitative approach of exploring brain functions such as mental chronometry and relative processing time, leading to a more elaborated description of cognitive models that include relevant timing information (Bellgowan et al., 2003). We hypothesize that early predictive activity in the occipital lobe prepares for the processing of the predicted visual feedback of self-generated as opposed to externally generated movements, leading to a reduced BOLD response amplitude and reduced BOLD duration when the predicted feedback is subsequently processed. This prediction related reduced processing goes along with reduced behavioral performance, in line with the cancelation account. An alternative explanation would be that the earlier processing leads to a temporally earlier representation of the feedback (feeling that the movement occurred earlier) compared to action, making the detection of small delays more difficult. This explanation could explain the often observed temporal binding effect (Hughes et al., 2013) by predictive shifts in the temporal representation of action outcomes.

### 4.4 Neuro-behavioral relationships

The timing of the hemodynamic response (TD) and behavioral threshold values in the left putamen, were correlated positively. That indicates that earlier activation in putamen in active versus passive conditions was linked with sensory suppression (worse performance in delay detection task). Bold amplitude suppression (passive>active) in bilateral precentral gyrus, in left middle occipital gyrus, lingual gyrus, superior frontal gyrus, putamen, rolandic operculum and in right superior occipital gyrus was correlated with worse performance in detecting delays (sensory attenuation) in self-generated versus externally generated movements. These correlations were not observed in our previous study (Arikan et al., 2019), so probably our results benefit from the use of basis function model, which is in agreement with a study reporting that the use of a better fitting model has a lot of advantages (Lindquist et al., 2009), such as the identification of meaningful brain-behavioral relationships. Shorter processing time (DD), in active as opposed to passive movements, was correlated with delay detection performance. Specifically, shorter processing time in left superior frontal gyrus, putamen, middle temporal gyrus, caudate, right cerebellum crus II, rolandic operculum, and in right SOG were correlated with worse performance in detecting delays (sensory attenuation) in self-generated versus externally generated movements.

### 4.5 Limitations

While the applied approach is suitable to examine whether forward model predictions lead to earlier and shorter processing of self-generated compared to external generated moments, this approach has some limitations. First, by identifying earlier processing with regard to the feedback movement onset it is now difficult to disentangle action and feedback related processes. Thus, early activity for example in the premotor cortex, SMA and cerebellum could be more related to action execution than the processing of the visual feedback. However, in this context we think it is impossible to disentangle action and related feedback processing anyways. At least the exploratory correlation analyses support our interpretation that areas with earlier processing are relevant for feedback processing. Secondly, we compared different regions regarding their individual timing which could be problematic due to differences in size, location regarding slice acquisition order and anatomical properties. However, as we always compare active and passive conditions within each regions, the relative timing might still be interpretable. Nevertheless, the development of new paradigms and data acquisition procedures as well as the application of more time sensitive methods are important to validate our results and to become a complete picture about the temporal sequence of involved processing steps.

### 4.6 Conclusion

To summarize, in line with our hypotheses we found not only reduced but also earlier and shorter BOLD responses in multiple cortical, subcortical and cerebellar brain structures. We found earlier BOLD activity in visual and somatosensory cortical, cerebellar and subcortical areas (basal ganglia and thalamus) during self- as opposed to externally generated movements, indicating that efference copy based predictive mechanisms enabled earlier processing of action feedback in self-generated movements. Also, we identified the sequencing of the neural activity during action feedback processing of self-generated movements, indicating earliest processing in occipital structures and the SMA, followed by predominantly cerebellar, parietal, and subcortical structures followed by sensory areas (postcentral gyrus). Finally, we reported for the first time, shorter BOLD duration in cortical and sub-cortical brain regions in self- versus externally generated movements, which was correlated with sensory attenuation in self-generated movements. Together, these new results highlight the relevance of considering timing and duration differences in BOLD responses when investigating action related predictive mechanisms. Furthermore, the data suggest that the consideration of timing and duration in addition to amplitude is important to predict and understand human behavior, which seem to be specifically related to the different aspects of the BOLD response. Finally, our results shed new light on the cortico-cerebellar-striatal loops involved during predictive perception of the visual feedback of one’s own hand movements. Together, these results provide a solid basis for investigating the role of BOLD timing and duration on impaired predictive mechanisms and action monitoring in patients with schizophrenia spectrum disorders (e.g., Blakemore et al., 2000; Leube et al., 2010; Uhlmann et al., 2021; Straube et al., 2020).

## Acknowledgements

This study was funded by the “Deutsche Forschungsgemeinschaft” (DFG) through the SFB/Transregio 135, “Cardinal mechanisms of perception: prediction, valuation, categorization”, and through the International Research Training Group, IRTG 1901, “The Brain in Action-BrainAct”) and the Hessisches Ministerium für Wissenschaft und Kunst (HMWK; project ‘The Adaptive Mind’). B.S. is supported by Deutsche Forschungsgemeinschaft grants STR 1146/8-1 and STR 1146/9-1. The authors confirm that there are no known conflicts of interest associated with this publication. We thank Jens Sommer for technical support and Christina Lubinus for assistance with data collection. The MRI data acquisition was supported by the Core Facility Brain Imaging, Faculty of Medicine, Philipps University Marburg, Germany.

## References

Alexander, G.E., DeLong, M.R., Strick, P.L., 1986. Parallel organization of functionally segregated circuits linking basal ganglia and cortex. Annu. Rev. Neurosci. VOL. 9, 357–381. https://doi.org/10.1146/annurev.ne.09.030186.002041

Arikan, B.E., van Kemenade, B.M.V., Podranski, K., Steinsträter, O., Straube, B., Kircher, T., 2019. Perceiving your hand moving: BOLD suppression in sensory cortices and the role of the cerebellum in the detection of feedback delays. J. Vis. 19, 1–22. https://doi.org/10.1167/19.14.4

Ball, T., Schreiber, A., Feige, B., Wagner, M., Lücking, C.H., Kristeva-Feige, R., 1999. The role of higher-order motor areas in voluntary movement as revealed by high-resolution EEG and fMRI. Neuroimage 10, 682–694. https://doi.org/10.1006/nimg.1999.0507

Bellgowan, P.S.F., Saad, Z.S., Bandettini, P.A., 2003. Understanding neural system dynamics through task modulation and measurement of functional MRI amplitude, latency, and width. Proc. Natl. Acad. Sci. U. S. A. 100, 1415–1419. https://doi.org/10.1073/pnas.0337747100

Blakemore, S.J., Frith, C.D., Wolpert, D.M., 1999. Spatio-temporal prediction modulates the perception of self-produced stimuli. J. Cogn. Neurosci. 11, 551–559. https://doi.org/10.1162/089892999563607

Blakemore, S.J., Sirigu, A., 2003. Action prediction in the cerebellum and in the parietal lobe, in: Experimental Brain Research. Exp Brain Res, pp. 239–245. https://doi.org/10.1007/s00221-003-1597-z

Blakemore, S.J., Wolpert, D.M., Frith, C.D., 1998. Central cancellation of self-produced tickle sensation. Nat. Neurosci. 1, 635–640. https://doi.org/10.1038/2870

Brass, M., Haggard, P., 2010. The hidden side of intentional action: the role of the anterior insular cortex. Brain Struct. Funct. https://doi.org/10.1007/s00429-010-0269-6

Calhoun, V.D., Stevens, M.C., Pearlson, G.D., Kiehl, K.A., 2004. fMRI analysis with the general linear model: Removal of latency-induced amplitude bias by incorporation of hemodynamic derivative terms. Neuroimage 22, 252–257. https://doi.org/10.1016/j.neuroimage.2003.12.029

Cignetti, F., Salvia, E., Anton, J.-L., Grosbras, M.-H., Assaiante, C., 2016. Pros and Cons of Using the Informed Basis Set to Account for Hemodynamic Response Variability with Developmental Data. Front. Neurosci. 10, 322. https://doi.org/10.3389/fnins.2016.00322

Crowell, R.M., Perret, E., Siegfried, J., Villoz, J.P., 1968. “Movement units” and “tremor phasic units” in the human thalamus. Brain Res. 11, 481–488. https://doi.org/10.1016/0006-8993(68)90142-X

Cui, R.Q., Deecke, L., 1999. High resolution DC-EEG analysis of the Bereitschaftspotential and post movement onset potentials accompanying uni-or bilateral voluntary finger movements. Brain Topogr. 11, 233–249. https://doi.org/10.1023/a:1022237929908

Cui, S.-Z., Li, E.-Z., Zang, Y.-F., Weng, X.-C., Ivry, R., Wang, J.-J., 2000. Both sides of human cerebellum involved in preparation and e… : NeuroReport [WWW Document]. URL https://journals.lww.com/neuroreport/Abstract/2000/11270/Both_sides_of_human_cerebellum_involved_in.49.aspx (accessed 5.16.21).

Cunnington, R., Iansek, R., Bradshaw, J.L., Phillips, J.G., 1995. Movement-related potentials in parkinson’s disease: Presence and predictability of temporal and spatial cues. Brain 118, 935–950. https://doi.org/10.1093/brain/118.4.935

Cunnington, R., Windischberger, C., Deecke, L., Moser, E., 2002. The preparation and execution of self-initiated and externally-triggered movement: A study of event-related fMRI. Neuroimage 15, 373–385. https://doi.org/10.1006/nimg.2001.0976

Daw, N.D., Niv, Y., Dayan, P., 2005. Uncertainty-based competition between prefrontal and dorsolateral striatal systems for behavioral control. Nat. Neurosci. 8, 1704–1711. https://doi.org/10.1038/nn1560

Dayan, P., 2009. Goal-directed control and its antipodes. Neural Networks 22, 213–219. https://doi.org/10.1016/j.neunet.2009.03.004

Deecke, L., Kornhuber, H.H., 1978. An electrical sign of participation of the mesial “supplementary” motor cortex in human voluntary finger movement. Brain Res. 159, 473–476. https://doi.org/10.1016/0006-8993(78)90561-9

Deecke, L., Scheid, P., Kornhuber, H.H., 1969. Distribution of readiness potential, pre-motion positivity, and motor potential of the human cerebral cortex preceding voluntary finger movements. Exp. Brain Res. 7, 158–168. https://doi.org/10.1007/BF00235441

Deiber, M.P., Ibanez, V., Sadato, N., Hallett, M., 1996. Cerebral structures participating in motor preparation in humans: A positron emission tomography study. J. Neurophysiol. 75, 233–247. https://doi.org/10.1152/jn.1996.75.1.233

Desantis, A., Haggard, P., 2016. How actions shape perception: learning action-outcome relations and predicting sensory outcomes promote audio-visual temporal binding. Sci. Rep. 6. https://doi.org/10.1038/srep39086

Díaz-Hernández, E., Contreras-López, R., Sánchez-Fuentes, A., Rodríguez-Sibrían, L., Ramírez-Jarquín, J.O., Tecuapetla, F., 2018. The Thalamostriatal Projections Contribute to the Initiation and Execution of a Sequence of Movements. Neuron 100, 739-752.e5. https://doi.org/10.1016/j.neuron.2018.09.052

Doya, K., 1999. What are the computations of the cerebellum, the basal ganglia and the cerebral cortex? Neural Networks 12, 961–974. https://doi.org/10.1016/S0893-6080(99)00046-5

Dudman, J.T., Krakauer, J.W., 2016. The basal ganglia: From motor commands to the control of vigor. Curr. Opin. Neurobiol. https://doi.org/10.1016/j.conb.2016.02.005

Eccles, J.C., 1982. The initiation of voluntary movements by the supplementary motor area. Arch. Psychiatr. Nervenkr. 231, 423–441. https://doi.org/10.1007/BF00342722

Elsinger, C.L., Harrington, D.L., Rao, S.M., 2006. From preparation to online control: Reappraisal of neural circuitry mediating internally generated and externally guided actions. Neuroimage 31, 1177–1187. https://doi.org/10.1016/j.neuroimage.2006.01.041

Fermin, A.S.R., Yoshida, T., Yoshimoto, J., Ito, M., Tanaka, S.C., Doya, K., 2016. Model-based action planning involves cortico-cerebellar and basal ganglia networks. Sci. Rep. 6, 1–14. https://doi.org/10.1038/srep31378

François-Brosseau, F.E., Martinu, K., Strafella, A.P., Petrides, M., Simard, F., Monchi, O., 2009. Basal ganglia and frontal involvement in self-generated and externally-triggered finger movements in the dominant and non-dominant hand. Eur. J. Neurosci. 29, 1277–1286. https://doi.org/10.1111/j.1460-9568.2009.06671.x

Gerloff, C., Altenmüller, E., Dichgans, J., 1996. Disintegration and reorganization of cortical motor processing in two patients with cerebellar stroke. Electroencephalogr. Clin. Neurophysiol. 98, 59–68. https://doi.org/10.1016/0013-4694(95)00204-9

Gläscher, J.P., O’Doherty, J.P., 2010. Model-based approaches to neuroimaging: Combining reinforcement learning theory with fMRI data. Wiley Interdiscip. Rev. Cogn. Sci. 1, 501–510. https://doi.org/10.1002/wcs.57

Haggard, P., Clark, S., Kalogeras, J., 2002. Voluntary action and conscious awareness. Nat. Neurosci. 5, 382–385. https://doi.org/10.1038/nn827

Henson, R.N.A., Price, C.J., Rugg, M.D., Turner, R., Friston, K.J., 2002. Detecting latency differences in event-related BOLD responses: Application to words versus nonwords and initial versus repeated face presentations. Neuroimage 15, 83–97. https://doi.org/10.1006/nimg.2001.0940

Hughes, G., Desantis, A., Waszak, F., 2013. Mechanisms of intentional binding and sensory attenuation: The role of temporal prediction, temporal control, identity prediction, and motor prediction. Psychol. Bull. 139, 133–151. https://doi.org/10.1037/a0028566

Ikeda, A., Shibasaki, H., Nagamine, T., Terada, K., Kaji, R., Fukuyama, H., Kimura, J., 1994. Dissociation between contingent negative variation and Bereitschaftspotential in a patient with cerebellar efferent lesion. Electroencephalogr. Clin. Neurophysiol. 90, 359–364. https://doi.org/10.1016/0013-4694(94)90051-5

Jahanshahi, M., Jenkins, I.H., Brown, R.G., Marsden, C.D., Passingham, R.E., Brooks, D.J., 1995. Self-initiated versus externally triggered movements: I. An investigation using measurement of regional cerebral blood flow with PET and movement-related potentials in normal and parkinson’s disease subjects. Brain 118, 913–933. https://doi.org/10.1093/brain/118.4.913

Jäncke, L., Shah, N.J., Peters, M., 2000. Cortical activations in primary and secondary motor areas for complex bimanual movements in professional pianists. Cogn. Brain Res. 10, 177–183. https://doi.org/10.1016/S0926-6410(00)00028-8

Jenkins, I.H., Jahanshahi, M., Jueptner, M., Passingham, R.E., Brooks, D.J., 2000. Self-initiated versus externally triggered movements. II. The effect of movement predictability on regional cerebral blood flow. Brain 123, 1216–1228. https://doi.org/10.1093/brain/123.6.1216

Klaus, A., Plenz, D., 2016. A Low-Correlation Resting State of the Striatum during Cortical Avalanches and Its Role in Movement Suppression. PLoS Biol. 14, 1002582. https://doi.org/10.1371/journal.pbio.1002582

Kornhuber, H.H., Deecke, L., 1965. Hirnpotentialänderungen bei Willkürbewegungen und passiven Bewegungen des Menschen: Bereitschaftspotential und reafferente Potentiale. Pflugers Arch. Gesamte Physiol. Menschen Tiere 284, 1–17. https://doi.org/10.1007/BF00412364

Lee, K.M., Chang, K.H., Roh, J.K., 1999. Subregions within the supplementary motor area activated at different stages of movement preparation and execution. Neuroimage 9, 117–123. https://doi.org/10.1006/nimg.1998.0393

Lindquist, M.A., Meng Loh, J., Atlas, L.Y., Wager, T.D., 2009. Modeling the hemodynamic response function in fMRI: efficiency, bias and mis-modeling. Neuroimage 45. https://doi.org/10.1016/j.neuroimage.2008.10.065

MacMillan, M.L., Dostrovsky, J.O., Lozano, A.M., Hutchison, W.D., 2004. Involvement of Human Thalamic Neurons in Internally and Externally Generated Movements. J. Neurophysiol. 91, 1085–1090. https://doi.org/10.1152/jn.00835.2003

Miall, R.C., Wolpert, D.M., 1996. Forward models for physiological motor control. Neural Networks 9, 1265–1279. https://doi.org/10.1016/S0893-6080(96)00035-4

Mohamed M.A., Yousem D.M., Tekes A., Browner N.M., Calhoun V.D., 2003. Timing of Cortical Activation: A Latency-Resolved Event-Related Functional MR Imaging Study | American Journal of Neuroradiology [WWW Document]. URL http://www.ajnr.org/content/24/10/1967 (accessed 5.24.21).

Monchi, O., Petrides, M., Strafella, A.P., Worsley, K.J., Doyon, J., 2006. Functional role of the basal ganglia in the planning and execution of actions. Ann. Neurol. 59, 257–264. https://doi.org/10.1002/ana.20742

Nieuwenhuis, S., Slagter, H.A., Von Alting Geusau, N.J., Heslenfeld, D.J., Holroyd, C.B., 2005. Knowing good from bad: Differential activation of human cortical areas by positive and negative outcomes. Eur. J. Neurosci. 21, 3161–3168. https://doi.org/10.1111/j.1460-9568.2005.04152.x

Oldenburg, I.A., Sabatini, B.L., 2015. Antagonistic but Not Symmetric Regulation of Primary Motor Cortex by Basal Ganglia Direct and Indirect Pathways. Neuron 86, 1174–1181. https://doi.org/10.1016/j.neuron.2015.05.008

Owen, A.M., 1997. MINI-REVIEW The Functional Organization of Working Memory Processes Within Human Lateral Frontal Cortex: The Contribution of Functional Neuroimaging, European Journal of Neuroscience.

Papa, S.M., Artieda, J., Obeso, J.A., 1991. Cortical activity preceding self-initiated and externally triggered voluntary movement. Mov. Disord. 6, 217–224. https://doi.org/10.1002/mds.870060305

Paradiso, G., Cunic, D., Saint-Cyr, J.A., Hoque, T., Lozano, A.M., Lang, A.E., Chen, R., 2004. Involvement of human thalamus in the preparation of self-paced movement. Brain 127, 2717–2731. https://doi.org/10.1093/brain/awh288

Pazen, M., Uhlmann, L., van Kemenade, B.M., Steinsträter, O., Straube, B., Kircher, T., 2020. Predictive perception of self-generated movements: Commonalities and differences in the neural processing of tool and hand actions. Neuroimage 206, 116309. https://doi.org/10.1016/j.neuroimage.2019.116309

Pernet, C.R., 2014. Misconceptions in the use of the General Linear Model applied to functional MRI: A tutorial for junior neuro-imagers. Front. Neurosci. https://doi.org/10.3389/fnins.2014.00001

Rao, S.M., 1997. Distributed neural systems underlying the timing of movements. J. Neurosci. 17, 5528–5535. https://doi.org/10.1523/jneurosci.17-14-05528.1997

Redgrave, P., Prescott, T.J., Gurney, K., 1999. The basal ganglia: A vertebrate solution to the selection problem? Neuroscience. https://doi.org/10.1016/S0306-4522(98)00319-4

Richter, W., Andersen, P.M., Georgopoulos, A.P., Kim, S.G., 1997. Sequential activity in human motor areas during a delayed cued finger movement task studied by time-resolved fMRI. Neuroreport 8, 1257–1261. https://doi.org/10.1097/00001756-199703240-00040

Romo, R., Schultz, W., 1992. Role of primate basal ganglia and frontal cortex in the internal generation of movements - III. Neuronal activity in the supplementary motor area. Exp. Brain Res. 91, 396–407. https://doi.org/10.1007/BF00227836

Sakata, H., Itoh, K., Suzuki, Y., Nakamura, K., Watanabe, M., Igarashi, H., Nakada, T., 2017. Slow accumulations of neural activities in multiple cortical regions precede self-initiation of movement: An event-related fMRI study. eNeuro 4. https://doi.org/10.1523/ENEURO.0183-17.2017

Sasaki, K., Jinnai, K., Gemba, H., Hashimoto, S., Mizuno, N., 1979. Projection of the Cerebellar Dentate Nucleus onto the Frontal Association Cortex in Monkeys, Brain Res.

Schmalenbach, S.B., Billino, J., Kircher, T., van Kemenade, B.M., Straube, B., 2017. Links between gestures and multisensory processing: Individual differences suggest a compensation mechanism. Front. Psychol. 8, 1828. https://doi.org/10.3389/fpsyg.2017.01828

Schmitter, C. V., Steinsträter, O., Kircher, T., van Kemenade, B.M., Straube, B., 2021. Commonalities and differences in predictive neural processing of discrete vs continuous action feedback. Neuroimage 229, 117745. https://doi.org/10.1016/j.neuroimage.2021.117745

Schultz, W., Romo, R., 1992. Role of primate basal ganglia and frontal cortex in the internal generation of movements - I. Preparatory activity in the anterior striatum. Exp. Brain Res. 91, 363–384. https://doi.org/10.1007/BF00227834

Shibasaki, H., Barrett, G., Neshige, R., Hirata, I., Tomoda, H., 1986. Volitional movement is not preceded by cortical slow negativity in cerebellar dentate lesion in man. Brain Res. 368, 361–365. https://doi.org/10.1016/0006-8993(86)90582-2

Shibasaki, H., Hallett, M., 2006. What is the Bereitschaftspotential? Clin. Neurophysiol. https://doi.org/10.1016/j.clinph.2006.04.025

Sommer, S., Pollmann, S., 2016. Putamen Activation Represents an Intrinsic Positive Prediction Error Signal for Visual Search in Repeated Configurations. Open Neuroimag. J. 10, 126–138. https://doi.org/10.2174/1874440001610010126

Soon, C.S., Brass, M., Heinze, H.J., Haynes, J.D., 2008. Unconscious determinants of free decisions in the human brain. Nat. Neurosci. 11, 543– 545. https://doi.org/10.1038/nn.2112

Soon, C.S., He, A.H., Bode, S., Haynes, J.D., 2013. Predicting free choices for abstract intentions. Proc. Natl. Acad. Sci. U. S. A. 110, 6217– 6222. https://doi.org/10.1073/pnas.1212218110

Sperry, R.W., 1950. Neural basis of the spontaneous optokinetic response produced by visual inversion. J. Comp. Physiol. Psychol. 43, 482–489. https://doi.org/10.1037/h0055479

Straube, B., Van Kemenade, B.M., Arikan, B.E., Fiehler, K., Leube, D.T., Harris, L.R., Kircher, T., 2017. Predicting the multisensory consequences of one’s own action: Bold suppression in auditory and visual cortices. PLoS One 12, e0169131. https://doi.org/10.1371/journal.pone.0169131

Thura, D., Cisek, P., 2017. The Basal Ganglia Do Not Select Reach Targets but Control the Urgency of Commitment. Neuron 95, 1160-1170.e5. https://doi.org/10.1016/j.neuron.2017.07.039

Uhlmann, L., Pazen, M., van Kemenade, B.M., Kircher, T., Straube, B., 2020. Neural Correlates of Self-other Distinction in Patients with Schizophrenia Spectrum Disorders: The Roles of Agency and Hand Identity. Schizophr. Bull. https://doi.org/10.1093/schbul/sbaa186

Ullsperger, M., 2006. Performance monitoring in neurological and psychiatric patients, in: International Journal of Psychophysiology. Elsevier, pp. 59–69. https://doi.org/10.1016/j.ijpsycho.2005.06.010

Van Donkelaar, P., Stein, J.F., Passingham, R.E., Miall, R.C., 1999. Neuronal activity in the primate motor thalamus during visually triggered and internally generated limb movements. J. Neurophysiol. 82, 934–945. https://doi.org/10.1152/jn.1999.82.2.934

van Kemenade, B.M., Arikan, B.E., Kircher, T., Straube, B., 2017. The angular gyrus is a supramodal comparator area in action–outcome monitoring. Brain Struct. Funct. 222, 3691–3703. https://doi.org/10.1007/s00429-017-1428-9

van Kemenade, B.M., Arikan, B.E., Kircher, T., Straube, B., 2016. Predicting the sensory consequences of one’s own action: First evidence for multisensory facilitation. Attention, Perception, Psychophys. 78, 2515–2526. https://doi.org/10.3758/s13414-016-1189-1

Van Kemenade, B.M., Arikan, B.E., Podranski, K., Steinsträter, O., Kircher, T., Straube, B., 2019. Distinct Roles for the Cerebellum, Angular Gyrus, and Middle Temporal Gyrus in Action-Feedback Monitoring. Cereb. Cortex 29, 1520–1531. https://doi.org/10.1093/cercor/bhy048

von Holst, E., Mittelstaedt, H., 1950. Das Reafferenzprinzip - Wechselwirkungen zwischen Zentralnervensystem und Peripherie. Naturwissenschaften 37, 464–476. https://doi.org/10.1007/BF00622503

Warner, W.A., Sanchez, R., Dawoodian, A., Li, E., Momand, J., 2013. NIH Public Access 80, 631–637. https://doi.org/10.1111/j.1747-0285.2012.01428.x.Identification

Weilke, F., Spiegel, S., Boecker, H., Von Einsiedel, H.G., Conrad, B., Schwaiger, M., Erhard, P., 2001. Time-resolved fMRI of activation patterns in M1 and SMA during complex voluntary movement. J. Neurophysiol. 85, 1858–1863. https://doi.org/10.1152/jn.2001.85.5.1858

Wessel, K., Zeffiro, T., Toro, C., Hallett, M., 1997. Self-paced versus metronome-paced finger movements: A positron emission tomography study. J. Neuroimaging 7, 145–151. https://doi.org/10.1111/jon199773145

Wiese, H., Stude, P., Nebel, K., De Greiff, A., Forsting, M., Diener, H.C., Keidel, M., 2004. Movement preparation in self-initiated versus externally triggered movements: An event-related fMRI-study. Neurosci. Lett. 371, 220–225. https://doi.org/10.1016/j.neulet.2004.08.078

Wildgruber, D., Erb, M., Klose, U., Grodd, W., 1997. Sequential activation of supplementary motor area and primary motor cortex during self-paced finger movement in human evaluated by functional MRI. Neurosci. Lett. 227, 161–164. https://doi.org/10.1016/S0304-3940(97)00329-7

Wolpert, D.M., Diedrichsen, J., Flanagan, J.R., 2011. Principles of sensorimotor learning. Nat. Rev. Neurosci. https://doi.org/10.1038/nrn3112

Wolpert, D.M., Flanagan, J.R., 2001. Motor prediction. Curr. Biol. https://doi.org/10.1016/s0960-9822(01)00432-8

Wolpert, D.M., Ghahramani, Z., 2000. Computational principles of movement neuroscience. Nat. Neurosci. 3, 1212–1217. https://doi.org/10.1038/81497

Worsley K.J., Marrett S., Neelin P., Vandal A.C., Frinston K.J., E.A.C., 1996. A unified statistical approach for determining significant signals in images of cerebral activation - Worsley - 1996 - Human Brain Mapping - Wiley Online Library [WWW Document]. URL https://onlinelibrary.wiley.com/doi/10.1002/(SICI)1097-0193(1996)4:1%3C58::AID-HBM4%3E3.0.CO;2-O (accessed 5.16.21).

Worsley, K.J., Evans, A.C., Marrett, S., Neelin, P., 1992. A three-dimensional statistical analysis for CBF activation studies in human brain. J. Cereb. Blood Flow Metab. 12, 900–918. https://doi.org/10.1038/jcbfm.1992.127

Wunderlich, K., Dayan, P., Dolan, R.J., 2012. Mapping value based planning and extensively trained choice in the human brain. Nat. Neurosci. 15, 786–791. https://doi.org/10.1038/nn.3068

