## Supplementary material for "The effect of self- vs. externally generated actions on timing, duration and amplitude of BOLD response for visual feedback processing"

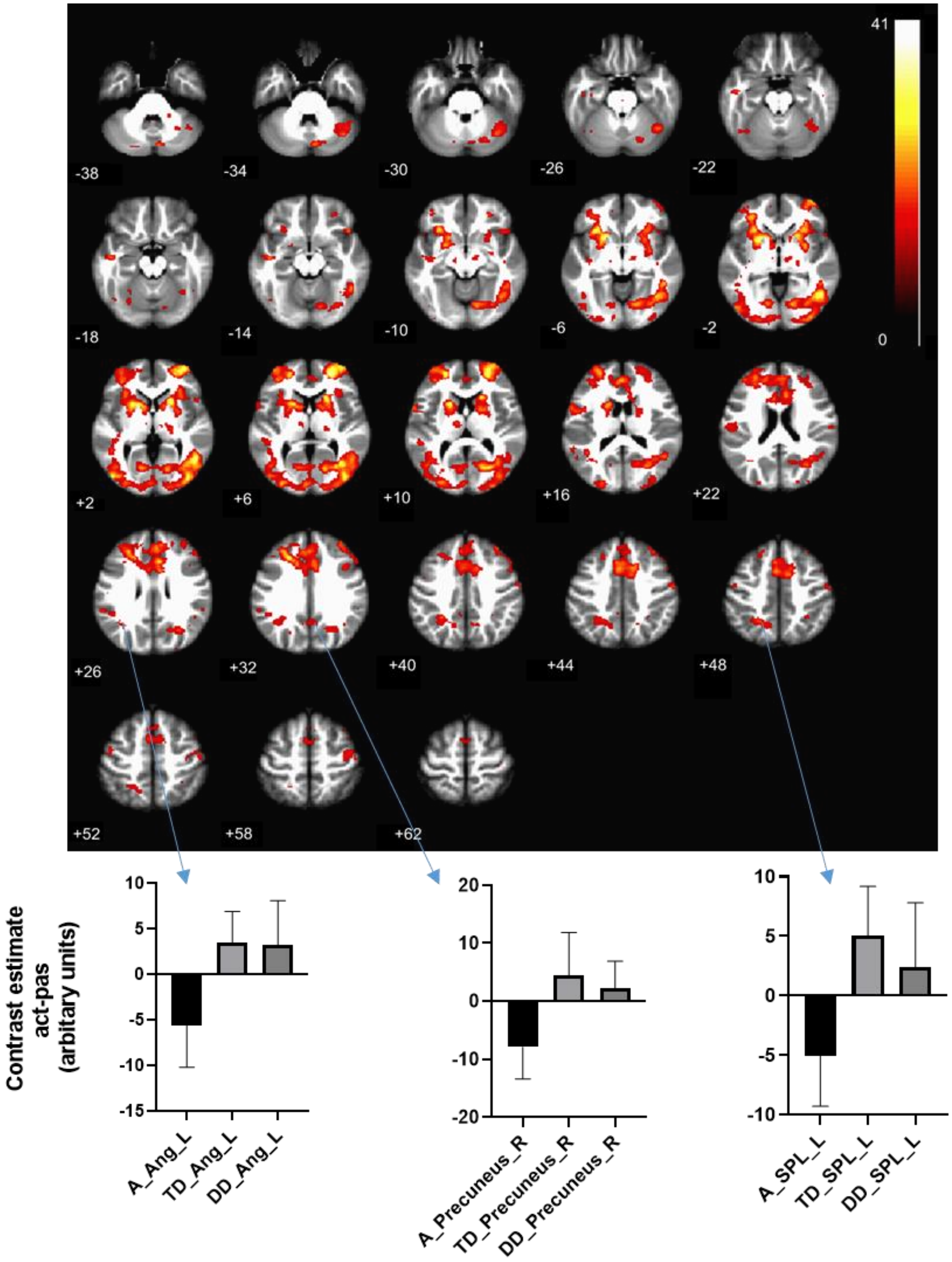

Figure 5. Clusters showing BOLD suppression, earlier activation and shorter BOLD duration in self-generated as opposed to externally generated movements. All maps were thresholded at pFWE < 0.05. A: Amplitude; TD: Temporal Derivative; DD: Dispersion Derivative. L: Left; R: Right. SPL: Superior Parietal Lobule; Ang: Angular gyrus.

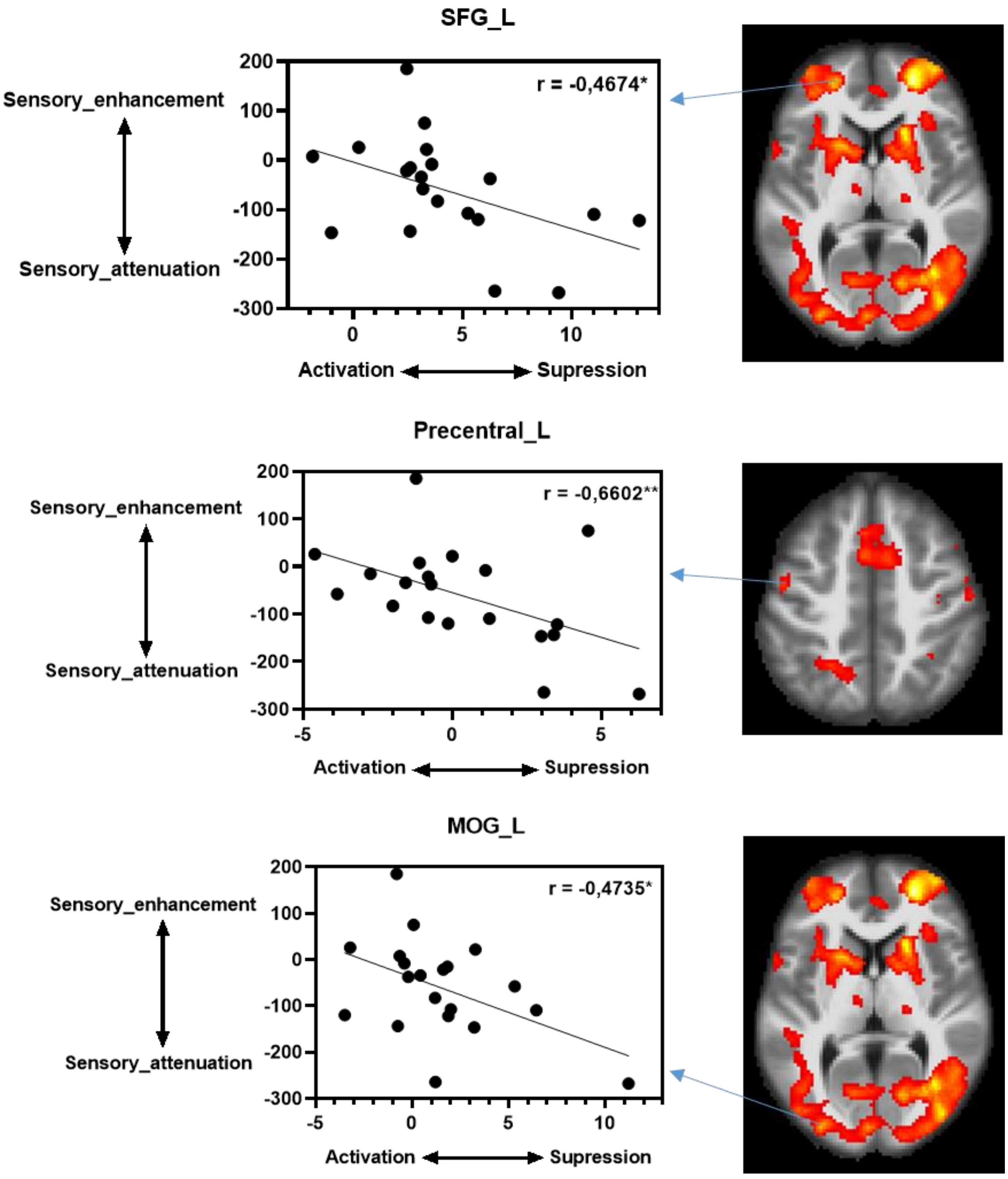

Figure 6. Pearson's correlation coefficient *r*, between BOLD amplitude values and threshold values of psychometric function from behavioral data (delay detection performance) in external generated minus self-generated movements (passive minus active contrast) are noted. Suppression in left superior frontal gyrus (SFG_L), in left precentral gyrus (Precentral_L) and in left middle occipital gyrus (MOG_L) was correlated with worse performance in detecting delays (sensory attenuation) in active versus passive conditions. The same trend was found in left lingual gyrus (r = -0.5316*), putamen (r = -0.4743*), rolandic operculum (r = -0.4571*) and in right precentral gyrus (r = -0.5080*), and right superior occipital gyrus (r = -0.4909*).

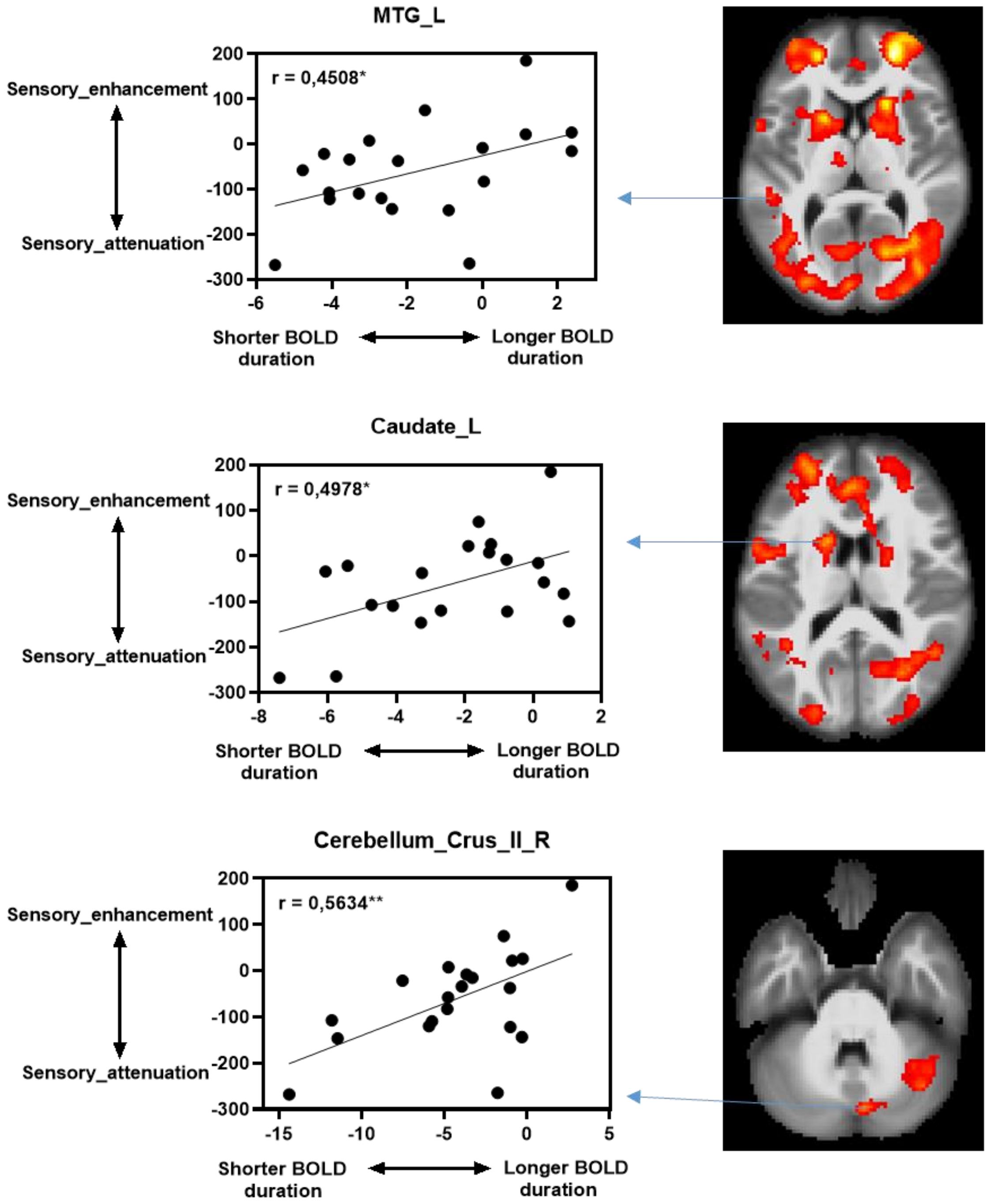

Figure 7. Pearson's correlation coefficient *r*, between BOLD processing time (Dispersion derivative estimates) and threshold values of psychometric function from behavioral data (delay detection performance) in external generated minus self-generated movements (passive minus active contrast). Shorter BOLD duration (processing time) in left middle temporal gyrus (MTG_L), in left Caudate and in right cerebellum crus II was correlated with worse performance in detecting delays (sensory attenuation) in active versus passive conditions. The same trend was found in left superior frontal gyrus ($r = 0.4676^*$), in left putamen ($r = 0.4416^*$), in left rolandic operculum ($r = 0.5128^*$), in left superior occipital gyrus ($r = 0.\ 4541^*$) and in right superior occipital gyrus ($r = 0.5008^*$).
